# Great-tailed Grackles (*Quiscalus mexicanus*) as a tolerant host of avian malaria parasites

**DOI:** 10.1101/2022.04.25.489425

**Authors:** M. Andreína Pacheco, Francisco C. Ferreira, Corina J. Logan, Kelsey B. McCune, Maggie P. MacPherson, Sergio Albino Miranda, Diego Santiago-Alarcon, Ananias A. Escalante

## Abstract

Great-tailed Grackles (*Quiscalus mexicanus*) are a social, polygamous bird species whose populations have rapidly expanded their geographic range across North America over the past century. Before 1865, Great-tailed Grackles were only documented in Central America, Mexico, and southern Texas in the USA. Given the rapid northern expansion of this species, it is relevant to study its role in the dynamics of avian blood parasites. Here, 87 Great-tailed grackles in Arizona (a population in the new center of the range) were screened for haemosporidian parasites using microscopy and PCR targeting the parasite mitochondrial cytochrome b gene. Individuals were caught in the wild from January 2018 until February 2020. Haemosporidian parasite prevalence was 60.9% (53/87). A high *Plasmodium* prevalence was found (59.8%, 52/87), and one grackle was infected with *Haemoproteus* (*Parahaemoproteus*) sp. (lineage SIAMEX01). Twenty-one grackles were infected with *P. cathemerium*, sixteen with *P. homopolare*, four with *P. relictum* (strain GRW04), and eleven with three different genetic lineages of *Plasmodium* spp. that have not been characterized to species level (MOLATE01, PHPAT01, and ZEMAC01). Gametocytes were observed in birds infected with three different *Plasmodium* lineages, revealing that grackles are competent hosts for some parasite species. This study also suggests that grackles are highly susceptible and develop chronic infections consistent with parasite tolerance, making them competent to transmit some generalist haemosporidian lineages. It can be hypothesized that, as the Great-tailed Grackle expands its geographic range, it may affect local bird communities by increasing the transmission of local parasites but not introducing new species into the parasite species pool.

## Introduction

The Great-tailed Grackle (*Quiscalus mexicanus*) is a sexually dimorphic, gregarious, and omnivorous passerine belonging to the family Icteridae. These social birds occur in North, Central, and South America, are polygamous, nest in colonies, and forage on the ground primarily in flocks from a few pairs to hundreds. They can be found in open and grassy areas with a source of surface water (e.g., pastures, agricultural areas, livestock feedlots, mangroves, secondary forests, second-growth scrub) and in urban landscapes (e.g., parks, garbage dumps, campuses) [1]. Such characteristics have allowed them to expand their geographic distribution by exploiting resources and habitats in human-modified environments, making them an urban dweller and utilizer species (*sensu* [2]).

The current distribution of the Great-tailed Grackle (henceforth grackles) ranges from the southern United States (mainly California, Arizona, New Mexico, and Texas) through Mexico and Central America to Colombia, Peru, and Venezuela [3]. Although this species is generally a short-distance and partial migrant in the northern parts of its range, it has recently become a permanent resident in areas where it formerly occurred only in the summer [1]. Indeed, its distribution has undergone a considerable, rapid, and large-scale expansion across the western United States in the last century. Grackles currently breed as far east as west Louisiana [4]. As the human population increased, changing the agricultural landscapes and urban centers of western USA, suitable habitats for grackles became more abundant and connected. This species was first reported in Arizona in 1935, where individuals were nesting in 1936 – 1937 before the agricultural boom in Sonora [5,6]. By 1964, the species became established in several places, and since then, their range has expanded northward [4].

Rapid range shifts can reveal plasticity in species’ traits. Such considerable change in the grackles range has driven studies focused on its systematics and population structure [7-14], behavior [15-19], reproduction and physiology [20-24], and geographic distribution and expansion [4,25-30]. However, only a few studies on this species have explored how it’s gregarious behavior and geographic expansion might affect local parasite communities and their biogeographical patterns [31]. Indeed, information on its parasites and pathogens is limited to a handful of studies on the West Nile Virus [32-35], *Toxoplasma gondii* [36], and hemoparasites [37].

Here, a haemosporidian parasite assemblage associated with this bird species in Arizona was studied as a first step to assessing its role in the parasite transmission dynamics. The potential role of this avian species as a competent and tolerant host of malaria parasites is also discussed. Avian haemosporidians are dipteran-borne parasites that belong to the genera *Plasmodium, Haemoproteus, Leucocytozoon*, and *Fallisia* [38]. Some of the species belonging to the genus *Plasmodium* are considered the most pathogenic and may cause malaria in their hosts; among them, *P. relictum* is globally widespread and has contributed to the extinction of many endemic species [39,40]. In this study, the haemosporidian parasite species were detected in a grackle population using microscopy and PCR targeting the parasite mitochondrial cytochrome b gene. This grackle population is considered one of the most recent in the middle of the northern expansion front.

## Materials and Methods

### Study area and samples

From January 2018 to April 2020, 87 grackles were caught in the wild in several locations around Tempe, a city in Maricopa County, Arizona, using walk-in traps, bow nets, and mist nets. The majority of individuals were caught on the campus of Arizona State University, while a small number were caught in a nearby city park. After capture, blood samples from each individual were obtained by brachial or medial metatarsal vein puncture, and then two to three thin blood smears were immediately prepared. The remaining blood sample was preserved in lysis buffer [41] for further molecular analysis. The birds were banded with colored leg bands in unique combinations for individual identification. Morphological measurements of weight, tarsus length, flattened wing length, tail length, skull length, bill length, and fat score (the amount of visible fat under the skin in the clavicle and abdomen as in [42]) were recorded. Such data was used to estimate the scaled mass index (SMI), which has become the primary method for quantifying energetic conditions within and among bird populations [43-45]. Then, grackles were either immediately released or temporarily brought into aviaries for behavioral testing (as part of other investigations [46-49]) and then released back to the wild. A second blood sample was collected and processed as described above from those grackles kept in the aviaries to confirm the haemosporidian diagnostic before being released. Grackles sampled twice were considered infected even if they were PCR-positive at only one of the time points.

### Examination of blood films and molecular detection of haemosporidian parasites

Blood smears were air-dried, fixed in absolute methanol for 5 min within 24 h of preparation, and then stained with 10% buffered Giemsa solution (pH 7.2) for 60 minutes. Blood films were examined with a Leica DM750 light microscope (Leica Microsystems, USA) equipped with a Leica ICC50 W camera and Leica Acquire software to capture images. Microscopic examination was carried out on blood smears that were deemed of high quality [50]. On those, 100 microscopic fields at low magnification (×400), and 200 fields at high magnification (×1000) were examined. Parasitemia (intensity of infection) was estimated as the number of parasites/20,000 erythrocytes counted at high magnification.

DNA from whole blood preserved in lysis buffer was extracted using QIAamp DNA Micro Kit (QIAGEN GmbH, Hilden, Germany) per the manufacturer’s protocol. Extracted DNA was screened to assess the presence of haemosporidians using a nested polymerase chain reaction (PCR) protocol that targets the parasite mitochondrial cytochrome b gene (*cytb*, 1,131 bp) with primers described by Pacheco et al. [51,52]. Primary PCR amplifications were carried out in a 50 µl volume with 4 µl of total genomic DNA, 2.5 mM MgCl2, 1 × PCR buffer, 0.25 mM of each deoxynucleoside triphosphate, 0.4 µM of each primer, and 0.03 U/µl AmpliTaq polymerase (Applied Biosystems, Thermo Fisher Scientific, USA). External PCR primers used were forward AE298 5′-TGT AAT GCC TAG ACG TAT TCC 3′ and reverse AE299 5′-GT CAA WCA AAC ATG AAT ATA GAC 3′. Primary PCR conditions were: A partial denaturation at 94 °C for 4 min and 36 cycles with 1 min at 94 °C, 1 min at 53 °C, and 2 min extension step at 72 °C. In the last cycle, a final extension step of 10 min at 72 °C was added. Nested PCR mix and conditions were the same as the primary PCR but using 1 µl of the primary PCRs and an annealing temperature of 56 °C. Internal PCR primers used were forward AE064 5′-T CTA TTA ATT TAG YWA AAG CAC 3′ and reverse AE066 5′-G CTT GGG AGC TGT AAT CAT AAT 3′. PCR amplified products (50 µl) were excised from agarose gels and purified by the QIAquick Gel Extraction Kit (QIAGEN GmbH, Hilden, Germany). Both strands for the *cytb* fragments were directly sequenced at Genewiz from Azenta Life Sciences (New Jersey, USA). It is worth noting that an advantage of the PCR protocol used here is that it yields richer phylogenetic information derived from a larger *cytb* fragment [51].

All electropherograms were carefully inspected, and samples with double peaks were considered as having either mixed infections (two genetic lineages of the same parasite species) or coinfections (two distinct parasite genera). Sequences obtained from mixed/coinfections were compared against lineages from the MalAvi database [53] and against those lineages identified in single infections in our samples. In addition, Poly Peak Parser online software was used to confirm lineage identity in mixed/coinfections [54]. All sequences obtained in this study were deposited in GenBank under the accession numbers ON227196-ON227265.

### Genetics and phylogenetic analyses

*Cytb* sequences obtained in this study were identified as *Plasmodium* or *Haemoproteus* using BLAST against GenBank [55] and MalAvi [53] databases. Once the lineage was identified, their prevalence in the grackle population was estimated and compared according to sex, weight, and body condition (SMI). Few individuals were caught per month (1-11) throughout the year, and only 6 in total during June, July, and August. Thus, the effect of seasonality on haemosporidian prevalence could not be determined.

Phylogenetic relationships between the lineages found in grackles and other haemosporidians were inferred using four alignments. These nucleotide alignments were performed using ClustalX v2.0.12 and Muscle as implemented in SeaView v4.3.543 [56]. The first alignment was constructed with 42 sequences and included 454 bp (excluding gaps) out of the 1,134 bp of *cytb* gene. This fragment is the most commonly sequenced *cytb* fragment [57] that allows broader comparisons between our samples and those deposited in public databases such as GenBank [58] and MalAvi [53]. This alignment included six *Plasmodium* lineages identified in this study, one *Plasmodium* lineage previously reported in a grackle from Arizona (KY653785, QUIMEX01), together with three *Plasmodium* sequences reported in grackles from Veracruz, Mexico (*Plasmodium homopolare* BAEBIC02, *Plasmodium nucleophilum* DENPET03, *Plasmodium* sp. QUIMEX02 [59]), as well as 28 *Plasmodium* identified to morphospecies level that had partial *cytb* gene data [60] available in GenBank [58] and MalAvi [53] at the time of this study. Four *Haemoproteus* (*Parahaemoproteus*) *cytb* sequences were included as an outgroup. A second alignment (N = 28) was done using the larger *cytb* fragment (945 bp out of the 1,134 bp, excluding gaps) [51] which overlaps with the most commonly sequenced *cytb* fragment [57]. This alignment included six *Plasmodium* lineages from this study, plus QUIMEX01 and those parasite species (N=22) with mtDNA genomes available [61]. Six *Haemoproteus* (*Parahaemoproteus*) and one *Leucocytozoon caulleryi* were included as an outgroup. Although it yielded a better phylogenetic signal than the phylogenetic tree using the small *cytb* fragment, it had less information in terms of isolates (N = 42 vs. 28). A third alignment (N=52) was done with the small *cytb* fragment sequences (467 out of the 1,134 bp of *cytb* gene, excluding gaps) of *Haemoproteus* (*Parahaemoproteus*) species (N=48) available in MalAvi database [53], including lineages from a previous study on grackles from Texas [37], and the *Haemoproteus* sp. sequence obtained in this study. In this case, four *Haemoproteus* (*Haemoproteus*) species were included as an outgroup. Finally, a fourth alignment (N = 22) was done using the larger *cytb* fragment (1,014 bp out of the 1,134 bp of *cytb* gene, excluding gaps) of those *Haemoproteus* sp. (N=20) with mtDNA genomes available [61], plus the *Haemoproteus* sp. lineage sequence obtained here. Four *Haemoproteus* (*Haemoproteus*) and one *Leucocytozoon caulleryi* were included as an outgroup.

Phylogenetic trees were estimated using the Bayesian method implemented in MrBayes v3.2.6 with the default priors [62], a general time-reversible model with gamma-distributed substitution rates, and a proportion of invariant sites (GTR + Γ + I). This model had the lowest Bayesian Information Criterion (BIC) scores, as estimated by MEGA v7.0.26 [63]. Bayesian support was inferred for the nodes in MrBayes by sampling every 1,000 generations from two independent chains lasting 2 × 10^6^ Markov Chain Monte Carlo (MCMC) steps. The chains were assumed to have converged once the potential scale reduction factor (PSRF) value was between 1.00 and 1.02, and the average SD of the posterior probability was < 0.01. Once convergence was reached, 25% of the samples were discarded as a “burn-in” [62]. GenBank accession numbers and the MalAvi lineage codes for all sequences used in the analyses are shown in the phylogenetic tree figures.

In addition, the first and third alignments were also used to estimate the pairwise evolutionary divergences among the genetic lineages of *Plasmodium* (N = 23, 454bp) and *Haemoproteus* species (N=5, 467bp) by using the Kimura 2-parameter model [64] as implemented in MEGA v7.0.26 [63].

### Statistical analyses

Test sensitivity was compared between PCR and microscopy using McNemar’s Chi-squared test with continuity correction in R (ver. 4.0.3; R Development Core Team 2020). Prevalence and their 95% confidence interval (CI) were estimated using the software Quantitative Parasitology on the Web (QPWeb) v.0.14 using the Clopper-Pearson method [65]. A Fisher exact test was used to determine non-random associations between being infected with *Plasmodium* and the host sex. In addition, a t-test was used to determine if there is a significant difference between infected and non-infected individuals in terms of (1) the mean of weight per sex and (2) SMI values estimated as in [66] per sex. These analyses were conducted during the first sampling before some of the birds were brought into the aviaries.

### Ethical statement and permits

This research was carried out following protocols and sample collection approved by the Institutional Animal Care and Use Committee at Arizona State University (protocol number 17-1594R) and under the permits from the US Fish and Wildlife Service (scientific collecting permit number MB76700A-0,1,2), US Geological Survey Bird Banding Laboratory (federal bird banding permit number 23872), and Arizona Game and Fish Department (scientific collecting license number SP594338 (2017), SP606267 (2018), SP639866 (2019), and SP402153 (2020)). All methods were performed following the relevant guidelines and regulations.

## Results

### Microscopic detection of haemosporidian parasites

Out of 87 sampled grackles, 69 high-quality blood smears were analyzed from a total of 55 birds, 14 of which had blood samples analyzed at two-time points. Forty-five samples were positive for haemosporidians by PCR, but parasites were detected by microscopy only in 17 blood smears. Ten of these samples had parasitemia (quantitative measure of parasites in the blood) ≤ 0.01%, and six samples had values between 0.015% and 0.02%. One grackle infected with the *Plasmodium* lineage PHPAT01 (identified by sequencing, see below) had parasitemia of 0.065% in the first sampling and a lower infection intensity four months later at the end of its time in the aviaries (parasitemia = 0.02%). Sequencing results from this second-time point revealed a different parasite (*P. homopolare*, LAIRI01), and this divergence in parasite identity was also confirmed morphologically (Fig 1A-E). From 24 PCR-negative samples, only one was positive by microscopy, displaying a 0.005% parasitemia.

**Figure 1.**
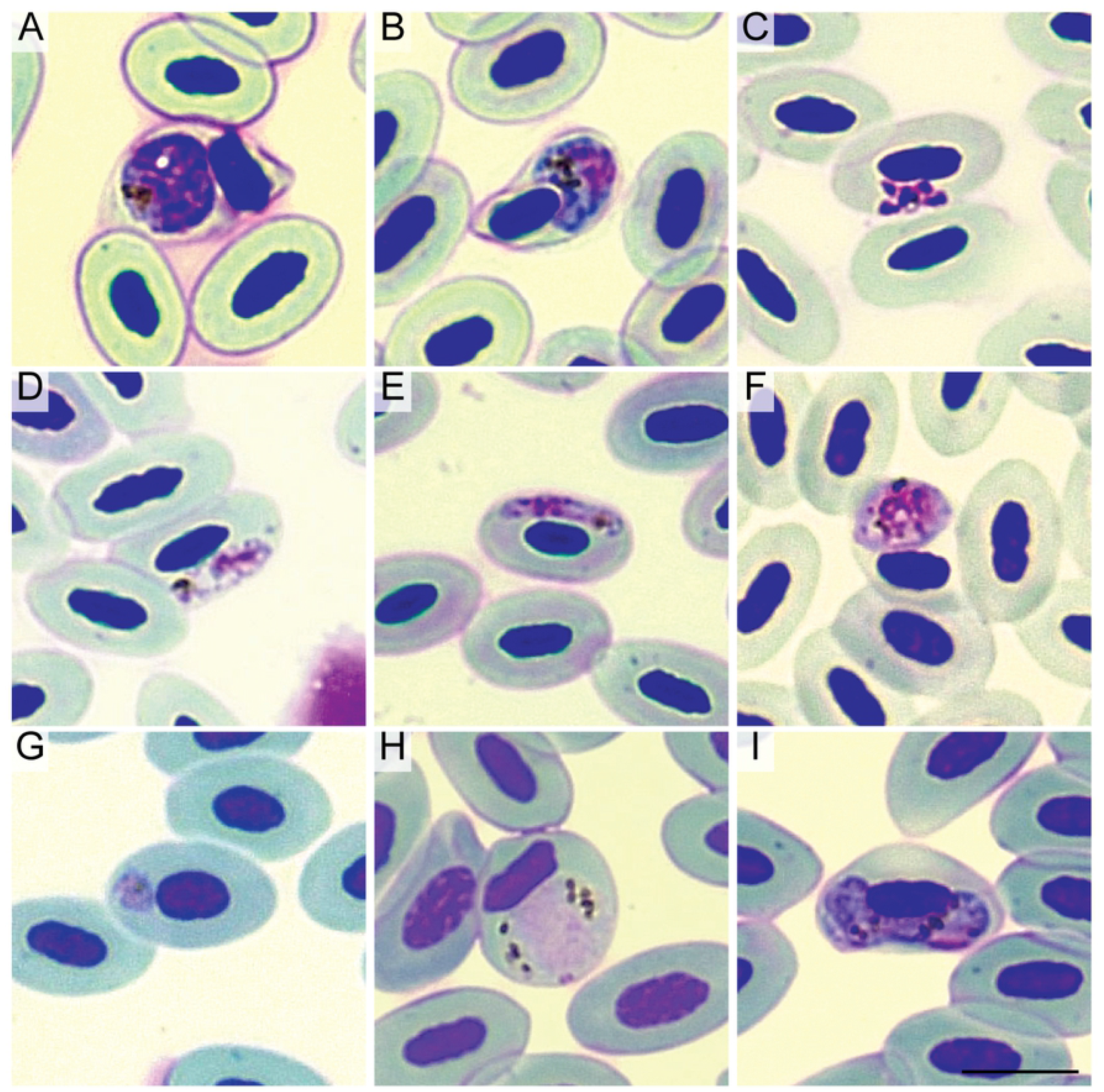
Blood stages of haemosporidian parasites infecting Great-tailed Grackles (*Quiscalus mexicanus*) from Tempe, Arizona. Meront (A) and macrogametocyte (B) visualized in a sample positive for *Plasmodium* sp. lineage PHPAT01. Meront (C) and gametocytes (D,E) of *Plasmodium homopolare* (LAIRI01). Gametocyte (F) in a sample positive for *Plasmodium* sp. lineage ZEMAC01. Immature parasite (G), likely a trophozoite, detected in a sample positive for *Plasmodium cathemerium*. Microgametocyte morphologically similar to *P. cathemerium* (H) and *Haemoproteus* sp. macrogametocyte (I) detected in a sample positive for *Haemoproteus* sp. (SIAMEX01). Giemsa-stained blood films. Scale bar = 10 μm.

*Plasmodium* gametocytes were detected in blood smears from samples that were PCR-positive for *Plasmodium* sp. (PHPAT01, Fig 1B), *P. homopolare* (LAIRI01, Fig 1D-E), and *Plasmodium* sp. (ZEMAC01, Fig 1F), confirming that grackles are competent hosts (the ability of a host to effectively transmit the parasite to a vector) for these parasites. No gametocytes were detected in samples positive for *Plasmodium* sp. (MOLATE01), *P. relictum* (GRW04), and *P. cathemerium* (SEIAUR01). However, immature parasites, likely to be trophozoites, were detected in 5 samples that were positive for the latter parasite (Fig 1G). One microgametocyte morphologically compatible with *P. cathemerium* (Fig 1H; [38]) was detected in a sample that was found infected with *Haemoproteus* by PCR with no signs of mixed infection (no double peaks visible in the electropherogram). A single *Haemoproteus* macrogametocyte was also detected in this blood smear (Fig 1I, parasitemia = 0,005%).

### Molecular detection of haemosporidian parasites, genetics, and phylogenetic analyses

Detectability of haemosporidians was strongly discordant between microscopy and PCR (McNemar’s Chi-squared = 23.31, df = 1, *p* < 0.0001), with a higher detection rate in the latter. The nested PCRs detected a high haemosporidian prevalence in this grackle population from Tempe, Arizona (60.9%, n = 87; Table 1). *Plasmodium* prevalence was higher (59.8%) compared to *Haemoproteus* (1.1%, Table 1). There was no association between being infected and the host sex (*p*-value = 0.714). Overall, having a *Plasmodium* infection was not associated with differences in body weight and SMI for males and females (*p* > 0.05).

**Table 1.**
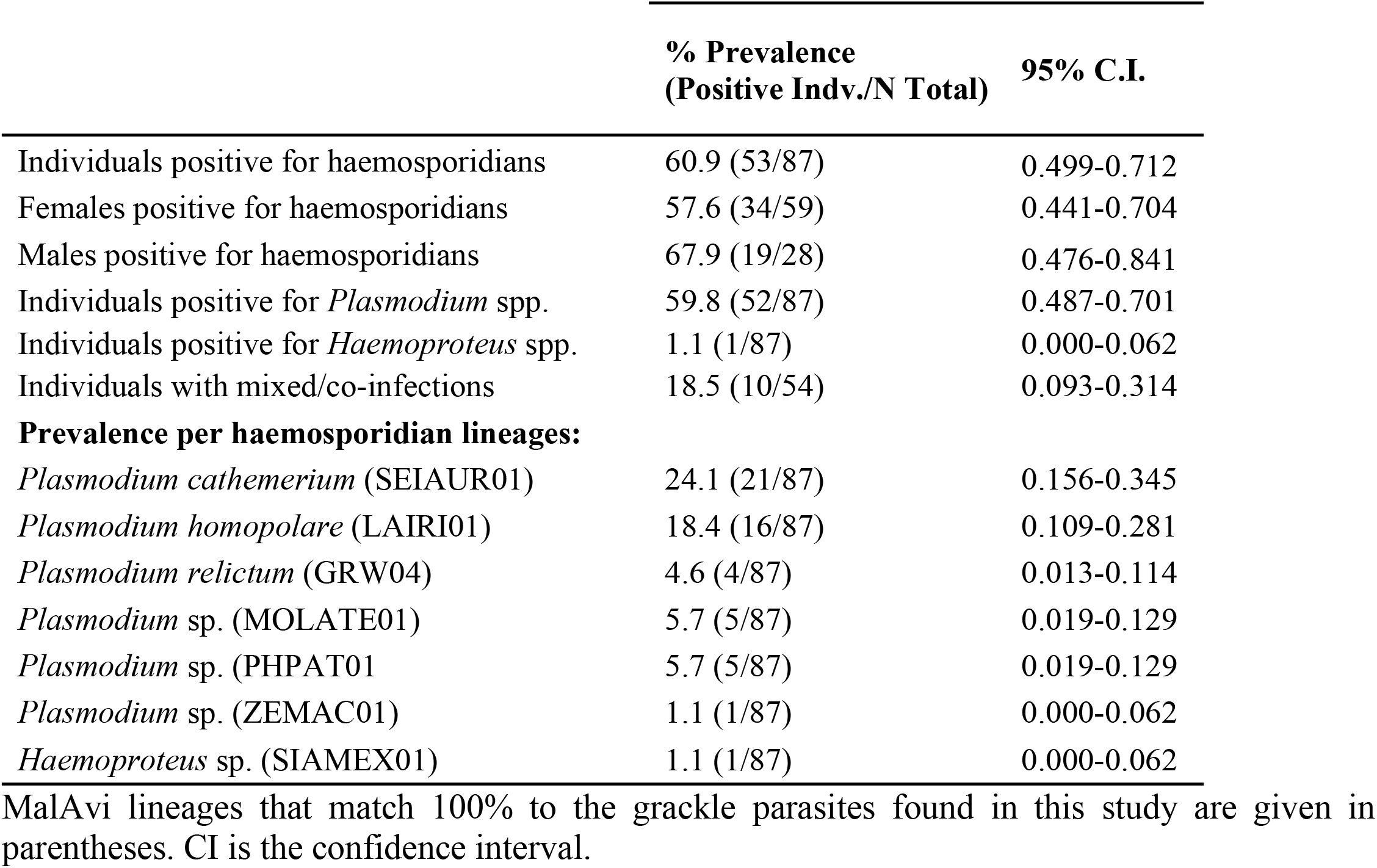
Prevalence of haemosporidian parasites in a Great-tailed Grackle (*Quiscalus mexicanus*) population sampled in Tempe, Arizona, between January 2018 and April 2020.

Ten grackles (18.5%) showed mixed infections (two different *Plasmodium* lineages) by PCR, and only one individual (1.9%) showed a coinfection of *Haemoproteus* and *Plasmodium* by microscopy. Six out of ten mixed infections were solved using Poly Peak Parser online software, and the combination of GRW04 and MOLATE01 was found in two individuals, SEIAUR01 and PHPAT01 in three, LAIRI01 and SEIAUR01 in one.

By comparing the sequences from grackles with single infections with those deposited in public databases [53,55], six *Plasmodium* lineages were detected (Tables 1). *Plasmodium cathemerium* (lineage SEIAUR01) and *P. homopolare* (lineage LAIRI01) were the most prevalent parasite species in this population (Tables 1 and 2, Fig 2). *Plasmodium relictum* lineage GRW04 was detected in low prevalence (Tables 1 and 2).

**Figure 2.**
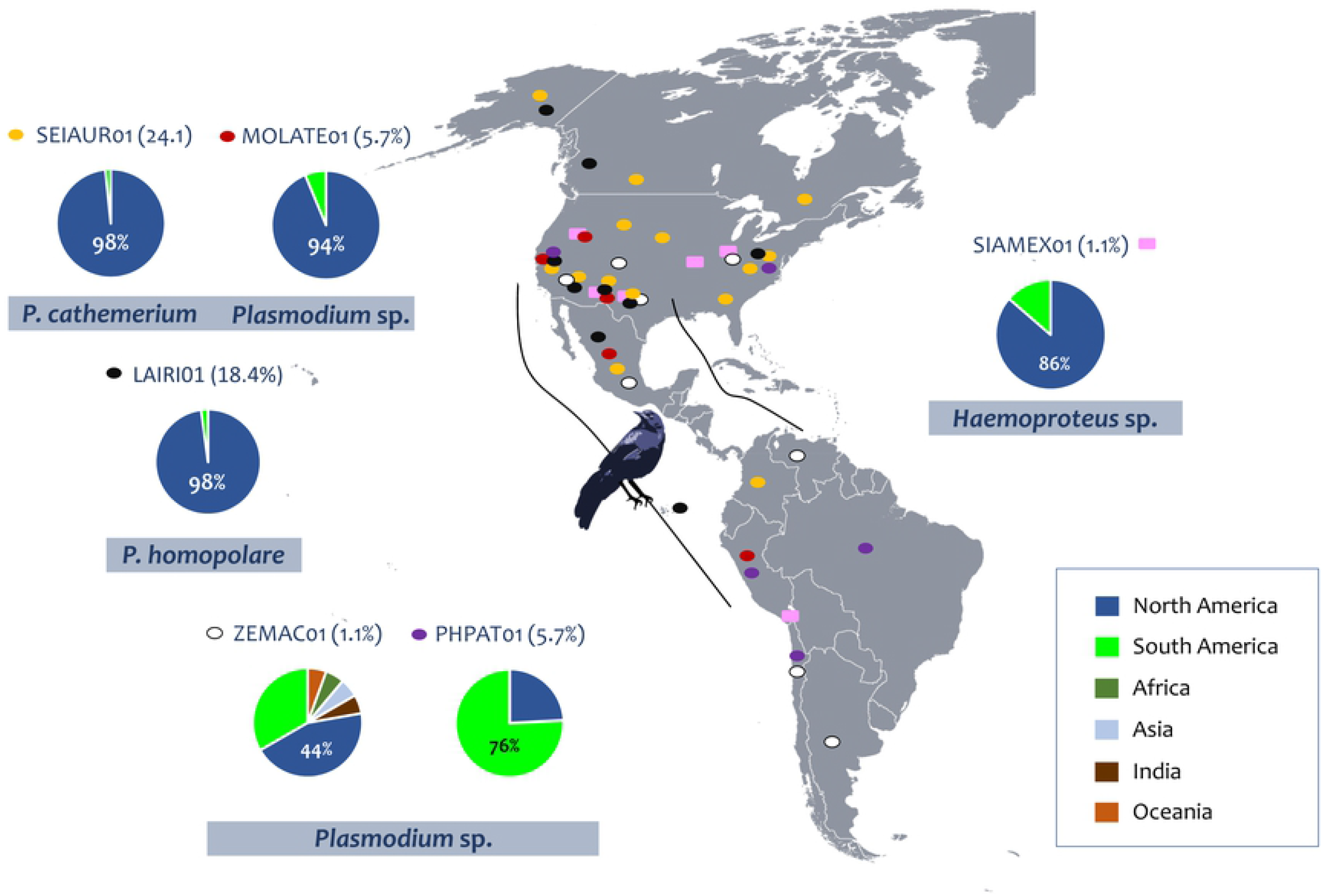
Map of the American continent showing the geographical distribution of haemosporidian lineages found in this study. The prevalence of each lineage is given in parenthesis. Color squares and values in the charts indicate continents and the global distribution of these lineages (MalAvi database, [53]), respectively, and color ovals represent the lineages. The distribution of *P. relictum* can be found in [40]. The grackle expansion range is indicated with black lines. Ariana Cristina Pacheco Negrin designed the map and grackle silhouette.

Genetic distance estimates among these six *cytb Plasmodium* lineages are shown in S1A Table. The genetic distance between *P. cathemerium* (lineage SEIAUR01) and the lineage MOLATE01 was low (0.009 ± 0.004; S1A Table) and comparable to the genetic variation between lineages of parasites belonging to the same morphospecies (*P. relictum* GRW04 vs. GRW11 = 0.020 ± 0.006; S1A Table). The genetic distance between lineages PHPAT01 and ZEMAC01 was lower (0.016 ± 0.006) than the genetic distance between two closely related *Plasmodium* species (e.g., *P. unalis* and *P. vaughani* = 0.033 ± 0.008; S1A Table). It was also low compared to the genetic distance between two lineages of *P. relictum* (e.g., GRW04 vs. GRW11 = 0.020 ± 0.006; S1A Table). In the case of *Haemoproteus*, the genetic lineage SIAMEX01 found in the Arizonan grackles was 100% identical (S1B Table) to a previously reported lineage SIAMEX01 found in Western bluebirds (*Sialia mexicana*) and lineage KF028P/CHI18PA found in grackles and the Tufted Titmouse (*Baeolophus bicolor*) from Texas [67].

The phylogenies obtained with the *Plasmodium* smaller (454 bp out of the 1,134 bp of *cytb* gene, excluding gaps) and larger *cytb* fragments (945 bp out of the 1,134 bp of *cytb* gene, excluding gaps) are shown in Fig 3 and S1A Fig respectively. The tree obtained using the larger fragment (S1A Fig) solved the relationship between the *Plasmodium* species with better support values in several clades, highlighting the importance of using longer *cytb* fragments when using phylogenetic methods. Both phylogenetic analyses share similarities and placed Arizonan grackle parasite sequences in four different clades.

**Figure. 3.**
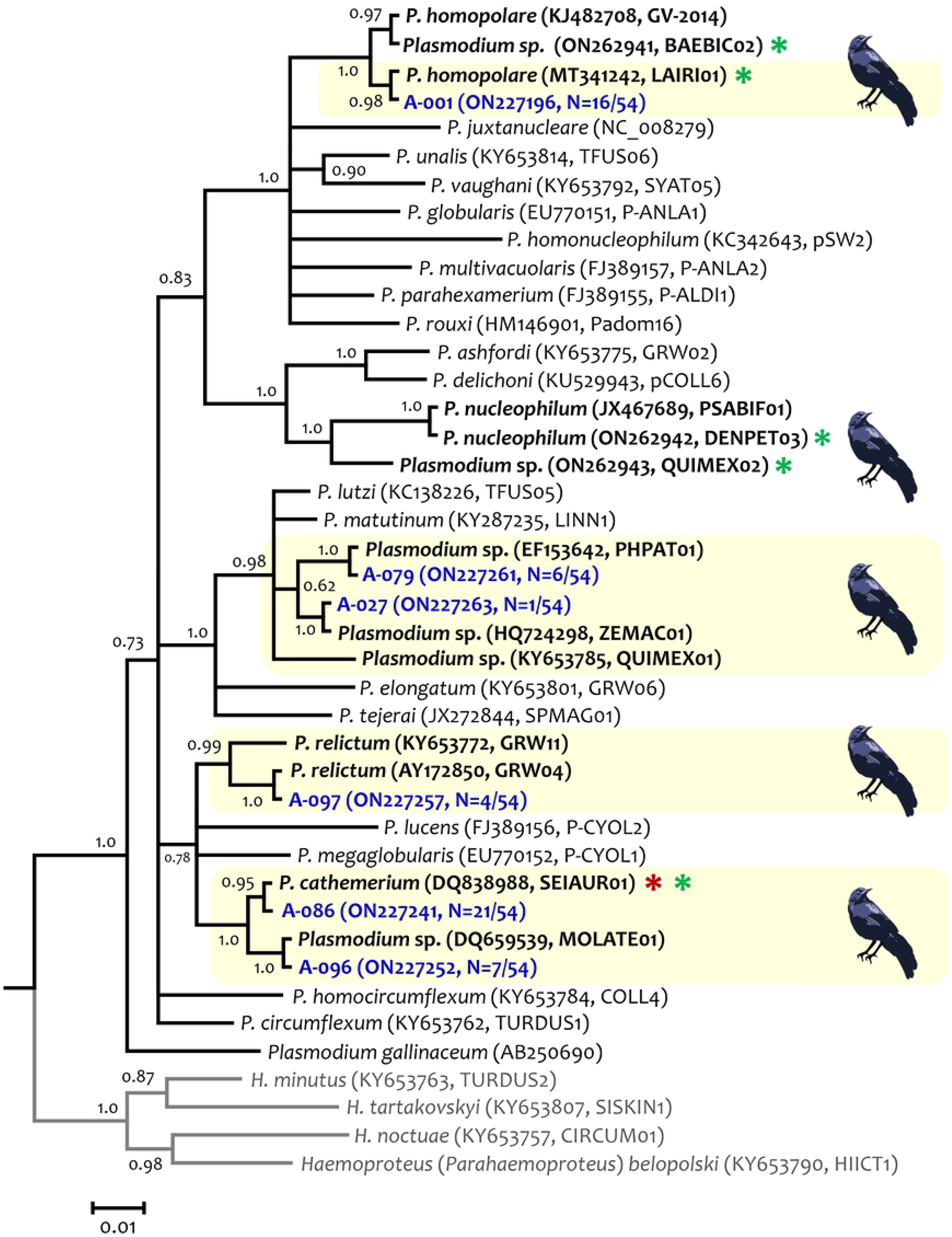
Bayesian phylogenetic hypothesis of *Plasmodium* parasites infecting Great-tailed Grackles (*Quiscalus mexicanus*) from Arizona, USA. Phylogenetic tree was computed based on parasites smaller *cytb* gene fragment (454 bp out of the 1,134 bp of *cytb* gene, excluding gaps). The values above branches are posterior probabilities. *Haemoproteus* (*Parahaemoproteus*) spp. (outgroup) are indicated in grey. Genbank accession numbers and their lineage identifiers, as deposited in the MalAvi database, are provided in parenthesis for the sequences used in the analysis. *Plasmodium* lineages recovered from grackles are written in blue, and the total of individuals infected with a given lineage in relation to the number of *Plasmodium*-positive samples is indicated. *Plasmodium cathemerium* was also detected in grackles from Texas [37] and is indicated with a red asterisk and parasites reported in Mexico are indicated with a green asterisk. Light yellow boxes indicate the lineages found in Arizona.

*Plasmodium homopolare* lineage LAIRI01 (Fig 3) circulating in the grackle population has been reported in California Condor populations from Arizona (H2, MT341242; [68]). As expected by their genetic distance, lineages PHPAT01 and ZEMAC01 are closely related and form a monophyletic group with lineage QUIMEX01 (or MAP01, KY653785) also reported in a grackle from Arizona [61], and with *Plasmodium lutzi* and *Plasmodium matutinum* (Fig 3 and S1A Fig). In this analysis, *P. cathemerium* (lineage SEIAUR01) and the lineage MOLATE01 shared a recent common ancestor with *P. relictum* (GRW11 and GRW04).

Phylogenetic relationships between *Haemoproteus* parasites are shown in Fig 4 and S1B Fig. In both phylogenies, *Haemoproteus* sp. SIAMEX01 (=KF028P=CHI18PA) found in grackles from Arizona shared a common ancestor with a lineage reported in grackles from Texas (PIPMAC01, [37]) and with *Haemoproteus lanii* (RB1). The genetic distance between *H. lanii* and the *Haemoproteus* lineage found in grackles from Arizona and Texas was 0.039 ± 0.008 (S1B Table).

**Figure. 4.**
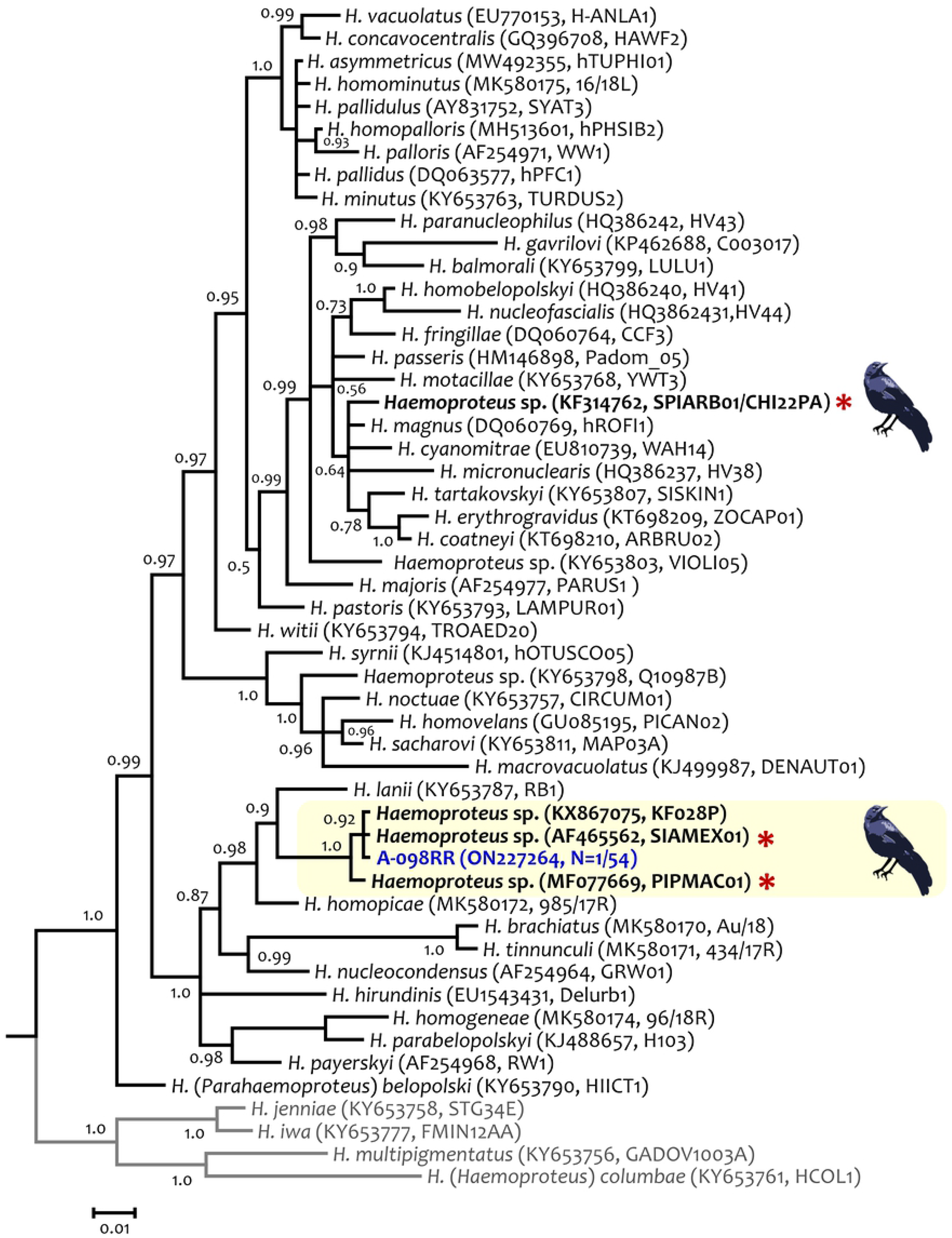
Bayesian phylogenetic hypothesis of *Haemoproteus* lineage infecting the Great-tailed Grackles (*Quiscalus mexicanus*) from Arizona, USA. The phylogenetic tree was computed based on parasites smaller *cytb* gene fragment (467 out of the 1,134 bp of *cytb* gene, excluding gaps). The values above branches are posterior probabilities. *Haemoproteus* (*Haemoproteus*) spp. (outgroup) are indicated in grey. Genbank accession numbers and lineage identifiers, as deposited in the MalAvi database, are provided in parenthesis for the sequences used in the analysis. *Haemoproteus* (*Parahaemoproteus*) lineage recovered from a grackle in this study is written in blue. Lineages detected in grackles from Texas [37] are indicated with a red asterisk.

## Discussion

In this study, a high haemosporidian prevalence (60.9%) and diversity of haemosporidian parasites (six *Plasmodium* lineages and one *Haemoproteus*) were detected in a grackle population from Arizona. This result contrasts with a previous study on 22 individuals in the same area that found no parasites using microscopy [69]. Such a difference could be explained, at least in part, by differences in sensitivity between PCR and microscopy in detecting haemosporidian parasites.

A prerequisite to making microscopy comparable with molecular detection methods is that experts extensively analyze high-quality blood smears [50], which was done in this study. However, no parasites were visualized in 60% of the positive samples by PCR. Indeed, parasitemias were below the detection level of standard microscopy (≤ 0.01% if only 10,000 erythrocytes are analyzed) in 59% of the blood smears in which parasites were detected. It has been shown that PCR can detect extremely low infection levels below the detectable microscopy threshold of approximately one parasite in a total of 50,000 examined cells [70]. This observation has been made in other Haemosporida, including human malarias [71]. A single parasite was detected in a blood smear from one PCR-negative sample out of 19 examined, revealing that the PCR assay used here is likely to produce few false-negative results. However, it is worth noting that microscopy allowed for detecting gametocytes (the infective parasite stage for vectors), evidence that grackles are competent hosts.

This study suggests that grackles are highly susceptible to infection. Also, the prevalent low parasitemias (undetectable by microscopy) and no signs of clinical disease in this population indicate chronically infected birds. Such characteristics are consistent with parasite tolerance [72], meaning that grackles could be competent reservoirs able to transmit some of the haemosporidian lineages found here.

There was no evidence of grackle-specific Haemosporida in this recent expansion front, and *Plasmodium* was the most common parasite (59.8 % prevalence) circulating in this population. All *Plasmodium* parasite lineages found here infect many residents and migrant species in the New World and are common in other bird species found in Arizona and other areas across the USA. These include lineages of *P. cathemerium, P. homopolare*, and *P. relictum* (S2 Table [38,40,68]). *Plasmodium cathemerium* (SEIUR01) has been detected in at least 33 bird species from 15 families (Passeriformes and non-Passeriformes), including House Sparrows (*Passer domesticus*, Passeridae) from Arizona [73], grackles from Mexico (S2 Table [59]) and other Icteridae species from California (Fig 2, S2 Table [53,55]). A *Plasmodium* sp. (MOLATE01, Table 1) related to *P. cathemerium* has been recovered from 9 bird species from 7 families – including Icteridae species (Passeriformes) – in the USA, Mexico, and Peru (Fig 2, S2 Table [53,55]).

*Plasmodium homopolare* (LAIRI01, Fig 2), on the other hand, has been reported in Masked Bobwhite Quails (*Colinus virginianus ridgwayi*, Odontophoridae), Common Ravens (*Corvus corax*, Corvidae) [74], and California Condors (*Gymnogyps californianus*, Cathartidae; [68]) from Arizona, and in grackles from Mexico (S2 Table [59]). In addition, the lineage LAIRI01 has been found in 26 bird species from 16 families (Passeriformes and non-Passeriformes), including other Icteridae species from nearby states (California and New Mexico) and across the Americas, with no reports in other continents (Fig 2, S2 Table [53,55]). Interestingly, Mexican grackles seem to be infected by the two *P. homopolare* lineages (LAIRI01 and BAEBIC02; [58]). In the case of *P. relictum*, the lineage GRW04 found in grackles is the most common *P. relictum* lineage in North America (>80%, [40]). The lineages found here in low prevalence, *Plasmodium* spp. ZEMAC01 (previously detected in 13 bird species from 12 families of Passeriformes and non-Passeriformes) and PHPAT01 (previously found in 26 bird species from 10 families of Passeriformes and non-Passeriformes) are also generalists (Fig 2, S2 Table [53,55,74]). As reported in Brazil, the lineage PHPAT01 can cause severe malaria in Magellanic Penguins [75]. Given that the genetic distance between PHPAT01 and ZEMAC01 is lower than the distance between two lineages of *P. relictum* (e.g., GRW04 vs. GRW11, S1A Table), it is worth exploring whether these are haplotypes of the same uncharacterized *Plasmodium* morphospecies.

The high *Plasmodium* prevalence in grackles found in Arizona contrasts with the results from a recent study showing that *Haemoproteus* (*Parahaemoproteus*) prevalence was considerably higher (73%, n = 44/60) than *Plasmodium* (1.7%, n = 1/60) in grackles sampled in Texas [37]. Out of the six *Plasmodium* genetic lineages detected in Arizona, only *P. cathemerium* (SEIAUR01, Figs 1 and 2) was found in a grackle from Texas (1/60). In contrast, out of the three *Haemoproteus* (*Parahaemoproteus*) lineages found in Texas (SPIARB01, SIAMEX01, and PIPMAC01), two (SIAMEX01, PIPMAC01) appear to be haplotypes of the only *Haemoproteus* sp. found in Arizona (SIAMEX01) (Figs 2 and 3, S1A Table). Interestingly, the only *Haemoproteus* detected in Arizona (SIAMEX01) was the most abundant lineage (68.6%) in the grackle population from Texas. This lineage seems to be also a generalist parasite as it has been previously recorded in 15 bird species from 10 families of Passeriformes and non-Passeriformes (S2 Table), including Icteridae species like Common Grackles from Michigan (*Quiscalus quiscula*, Icteridae; S2 Table S2) and Montezuma Oropendolas from Texas (*Psarocolius montezuma*). The life cycle of this *Haemoproteus* lineage and its pathology in different hosts remain unknown.

Although different sets of primers [76] were used for the nested PCRs in the study from Texas, all protocols employed here and in Golnar et al. [37] generally yield high sensitivity regardless of laboratory practice, parasite identity, and parasitemia [51,52,77]. *Haemoproteus* species were not found in blood smears from samples that were negative according to the PCR method, or in samples positive for *Plasmodium*, meaning that the PCR protocol employed here did not fail to detect *Haemoproteus* in our samples. Therefore, differences in parasite composition between grackles from Arizona (mostly infected by *Plasmodium*) and Texas (mostly infected by *Haemoproteus*) cannot be explained by methodological differences. It is worth noting that the only data available for haemosporidians in this species of grackles was the one from Texas [37] and some sequences available from Mexico [59]. Although no population studies on grackles have been conducted in Mexico, results obtained using similar protocols [59] from nine samples are consistent with the findings from Arizona. In particular, only *Plasmodium* spp. has been found, and four different parasite species (and five lineages) were detected by nested PCR (Fig 3; S2 Table [59]) in grackles sampled in Mexico.

The lineages found in grackles also can infect the two most common, year-round, and widespread birds, House Sparrows (an invasive species) and House Finches. These three species (grackles, House Sparrows, and House Finches) are found in sympatry in the study area (McCune KB, Personal Communication; https://ebird.org/barchart?byr=2018&eyr=2020&bmo=1&emo=12&r=US-AZ-013). The extent to which the transmission dynamics of haemosporidian parasites can be affected by tolerant and highly competent host species, or a guild of such host species, should be explored. Indeed, putative guilds of competent and relatively abundant hosts, including recently established species like the grackles or invasive species, such as the House Sparrow, are commonly found in urban and peri-urban areas. Specific inferences could be made if such a guild is relevant in terms of the haemosporidian parasite transmission in a particular area; for example, it could be expected that commonly abundant birds could favor some lineages of the local pool of parasite species. Such dynamics can affect the viability of other host species that exhibit less tolerance to certain parasites [74] or affect the composition of the local parasite species pool. Therefore, as the data suggests, it can be hypothesized that, as grackles expand geographically, they can transmit local generalist parasites rather than introducing new species into the parasite species pool. Thus, as a susceptible and tolerant host species [72], grackles could disproportionally transmit generalist parasites.

Avian *Plasmodium* species are transmitted by Culicidae insects, more commonly by *Culex* mosquitoes, while *Haemoproteus* (*Parahaemoproteus*) parasites are transmitted by biting midges of the genus *Culicoides* [78]. Grackles in the Arizona study area are found in urban landscapes with a source of surface water (e.g., ponds and lakes). Such sites are important for mosquito reproduction, increasing the probability of *Plasmodium* infections and transmission in the vicinity of lakes and ponds [79-82]. Also, it has been reported that *Plasmodium* is more prevalent than *Haemoproteus* species in urban greenspaces and peri-urban areas, suggesting that some parasite lineages could benefit from urbanization [83], which may explain the higher *Plasmodium* prevalence in Arizona. However, the urban area where grackles were sampled in Texas [37] has a high diversity of biting midges, and their molecular testing revealed a 2% infection rate for *Haemoproteus* in these insects [84]. It is possible that local differences in the vegetation and grackle behavior may expose them to different vectors. Bird species foraging at the canopy level have a higher probability of being infected with *Haemoproteus* than with *Plasmodium* [85], meaning that differences in foraging behavior between grackle populations could also contribute to contrasting results in haemosporidian assemblages. Thus, microclimatic conditions, vegetation, and differences in bird behavior may explain the differences between the populations from Texas and Arizona. Such differences may relate to land-use types and the specific environmental requirements of their different vectors, determining the local higher prevalence of one parasite genus over another.

In conclusion, the grackles are a highly susceptible and tolerant host species for haemosporidian parasites that harbor infections with low parasitemia. This species seems to acquire the local parasite species pool rather than expand the geographic range of particular “grackle-specific” lineages. Together with other species with similar characteristics, grackles may be part of community-level processes driving the parasite transmission of some haemosporidian lineages that remain poorly understood. Such dynamics involving guilds of multiple hosts remain unknown. It can be proposed that the haemosporidian species pool in bird assemblages containing guilds of tolerant and competent hosts in urban and peri-urban areas could be a suitable model for understanding the effects of host competence in a community context.

## Acknowledgments

The authors express their sincere gratitude to Melissa Folsom, Carolyn Rowney, and Luisa M. Bergeron for trapping and/or collecting/processing blood from some of the grackles involved in this research, and to Richard McElreath in the Department of Human Behavior, Ecology and Culture at the Max Planck Institute for Evolutionary Anthropology for funding. We thank Andrew J. Golnar and Gabriel L. Hamer for sharing the data obtained from Texas. We thank Ariana Cristina Pacheco Negrin for designing the map and grackle silhouette. FCF was supported by National Science Foundation (DEB 1717498). DS-A and SAM were supported by CONACYT, project number Ciencia Básica 2011-01-168524.

## Data Availability Statement

All sequences obtained in this study were deposited in GenBank under the accession numbers ON227196-ON227265.

## Author Contributions

**Conceptualization:** M. Andreína Pacheco, Ananias A. Escalante, Corina J. Logan, Francisco C. Ferreira.

**Data curation:** M. Andreína Pacheco.

**Formal analysis:** M. Andreína Pacheco, Francisco C. Ferreira.

**Funding acquisition:** Corina J. Logan.

**Investigation:** M. Andreína Pacheco, Francisco C. Ferreira, Corina J. Logan, Kelsey B. McCune, Sergio Albino Miranda, Diego Santiago-Alarcon and Ananias A. Escalante, Maggie

P. MacPherson..

**Methodology:** M. Andreína Pacheco, Francisco C. Ferreira, Corina J. Logan, Kelsey B. McCune, Sergio Albino Miranda.

**Project administration:** Corina J. Logan.

**Resources:** Corina J. Logan, Ananias A. Escalante.

**Supervision:** M. Andreína Pacheco, Ananias A. Escalante, Corina J. Logan, Kelsey B. McCune.

**Validation:** M. Andreína Pacheco, Francisco C. Ferreira, Ananias A. Escalante, Corina J. Logan.

**Visualization:** M. Andreína Pacheco, Francisco C. Ferreira.

**Writing – original draft:** M. Andreína Pacheco, Ananias A. Escalante.

**Writing – review & editing:** M. Andreína Pacheco, Ananias A. Escalante, Francisco C. Ferreira, Corina J. Logan, Sergio Albino Miranda, Diego Santiago-Alarcon, Kelsey B. McCune, Maggie P. MacPherson.

## Supporting information

### Supporting figures

**S1 Figure. Bayesian phylogenetic hypotheses of haemosporidian parasites infecting Great-tailed Grackles (*Quiscalus mexicanus*) from Arizona, USA**. Phylogenetic trees were computed based on larger fragments of (A) *Plasmodium* sequences of *cytb* gene (945 bp out of the 1,134 bp of *cytb* gene, excluding gaps) and (B) *Haemoproteus* sequences of *cytb* gene (1,014 bp out of the 1,134 bp of *cytb* gene, excluding gaps). Values above branches are posterior probabilities. *Leucocytozoon* and *Haemoproteus* (*Haemoproteus*) genera (outgroup) are indicated in grey. Genbank accession numbers and lineage identifiers, as deposited in the MalAvi database, are provided in parenthesis for the sequences used in the analyses. *Plasmodium* and *Haemoproteus* (*Parahaemoproteus*) recovered from grackles from Arizona are written in blue. Lineages detected in grackles from Texas [37] are indicated with a red asterisk and from Mexico with a green asterisk. **Supporting tables**.

**S1 Table. Estimates of evolutionary divergence between *cytb* (A) *Plasmodium* and (B)**

***Haemoproteus* parasites sequences**.

**S2 Table. GenBank and MalAvi records related to the haemosporidian lineages recovered from grackles from Arizona**. Bird species of the Icteridae family infected with the lineages found in this study are highlighted in blue and bird species from Arizona in yellow. The distribution and frequency of *P. relictum* can be found in [40].

